# Detrimental effects of advanced glycation end-products (AGEs) on a human neuromuscular junction co-culture model

**DOI:** 10.64898/2026.07.07.736594

**Authors:** Razan Alomosh, Amy Bateman, Kamel Mamchaoui, Vincent Mouly, Adam P. Lightfoot, Nessar Ahmed, Moi Hoon Yap, Nasser Al-Shanti

## Abstract

The neuromuscular junction (NMJ) is a specialised synapse between motor neurons and skeletal muscle, and its progressive deterioration contributes to age-related and metabolic disease-associated declines in muscle function. Advanced glycation end-products (AGEs) accumulate in tissues during ageing, diabetes, and chronic metabolic dysfunction and have been implicated in neuromuscular degeneration, yet their effects on the intact NMJ have not previously been examined in a human model system. This study employed a fully human, serum-free, and neural growth factor-free NMJ co-culture system, combining neural progenitor cells with immortalised human myoblasts derived from an 83-year-old donor, to investigate the effects of AGE exposure on neuromuscular integrity across structural, metabolic, functional, and secretory outcomes. AGE exposure induced significant reductions in motor neuron axonal length, myotube remodelling with centralised nuclear positioning, mitochondrial membrane depolarisation, elevated mitochondrial superoxide production, mitochondrial uncoupling, and reductions in spontaneous contraction intensity and frequency. Neurotrophic and myogenic growth factor signalling was significantly downregulated in AGE-treated co-cultures. These findings identify the NMJ as a sensitive target of glycation stress and establish this fully human co-culture platform as a physiologically relevant model for investigating glycation-related neuromuscular pathology and evaluating candidate therapeutic interventions.

## 1. Introduction

The neuromuscular junction (NMJ) is a specialised synapse between motor neurons and skeletal muscle fibres, through which precise neural control of muscle contraction is maintained. Progressive deterioration of neuromuscular connectivity is recognised as a contributing factor to age-related and disease-associated declines in muscle performance (Li, Xiong, & Mei, 2018; Pratt, De Vito, Narici, & Boreham, 2020).

Advanced glycation end-products (AGEs) are a heterogeneous group of bioactive compounds formed through the non-enzymatic glycation of proteins, lipids, and nucleic acids by reducing sugars, a process collectively termed the Maillard reaction (Zgutka, Tkacz, Tomasiak, & Tarnowski, 2023). AGEs accumulate progressively in tissues with age, metabolic dysfunction, and dietary intake, and their accumulation is markedly accelerated in conditions such as type 2 diabetes and chronic kidney disease. In skeletal muscle, AGEs promote oxidative stress, inflammation, and cross-linking of structural proteins, impairing tissue elasticity, cellular signalling, and contractile performance (Olson, Redden, Schwartz, Cohen, & McClure, 2021). These changes are particularly relevant to sarcopenia, the age-related loss of skeletal muscle mass, strength, and function, a condition increasingly recognised as a neuromuscular syndrome rather than a purely myogenic one (Gonzalez-Freire, de Cabo, Studenski, & Ferrucci, 2014). Elevated AGE levels have been associated with reduced muscle strength and functional decline in ageing populations, supporting the hypothesis that glycation stress contributes to neuromuscular degeneration (Guo et al., 2025).

The detrimental effects of AGEs on skeletal muscle have been investigated in both animal models and conventional two-dimensional (2D) cell culture systems (Adachi et al., 2019; Chiu et al., 2016; Zhao, Iyama, Kurogi, Hayashi, & Egawa, 2024). Whilst these studies have provided mechanistic insights, 2D systems and rodent models have limited translational relevance. More recent 3D microphysiological models have enabled physiologically meaningful assessments of AGE-induced damage (Kim et al., 2025), but have focused on muscle in isolation, without incorporating the motor neuron component integral to neuromuscular function.

Mitochondria localise specifically to the presynaptic terminal and junctional folds of the NMJ, where they not only sustain the energetic demands of synaptic transmission and contractile activity but also buffer the calcium ion loads essential for excitation-contraction coupling (Yipeng et al., 2025). In the context of sarcopenia, mitochondrial dysfunction is increasingly recognised as an early and potentially causal event in NMJ deterioration, preceding overt structural denervation (Chai et al., 2026; Miao et al., 2024). AGEs impair mitochondrial function in skeletal muscle through RAGE-dependent ROS generation, promoting membrane depolarisation, disruption of oxidative phosphorylation, and suppression of mitochondrial biogenesis via PGC-1α downregulation (Guo et al., 2025). Whether AGE-induced mitochondrial impairment specifically contributes to NMJ dysfunction in a human model system has not been previously examined.

The NMJ is a bidirectional signalling unit in which motor neurons and muscle fibres maintain continuous trophic and functional crosstalk (J. Saini et al., 2021). This microenvironment is enriched with neurotrophic factors, including BDNF, GDNF, and NT3, which are endogenously secreted to preserve axonal integrity, synaptic stability, and muscle fibre homeostasis (Davis, Fogarty, Brown, & Sieck, 2022). Whether AGE-induced glycation stress disrupts this neuromuscular signalling axis has not, to our knowledge, been investigated in a human in vitro system.

In the present study, we employed a fully human, serum-free, and neural growth factor-free NMJ co-culture system combining neural progenitor cells derived from the shef3 human embryonic stem cell line with immortalised human myoblasts (C83 cell line, derived from an elderly donor) to investigate the effects of AGEs on neuromuscular integrity. This system enables simultaneous assessment of neuronal and muscular responses to glycation stress within a functional NMJ microenvironment, preserving the bidirectional crosstalk between motor neurons and muscle that is absent in single-cell-type models. We hypothesised that AGE exposure would impair NMJ integrity across structural, metabolic, and functional levels, and that these effects would be reflected in the disruption of the trophic signalling environment.

## 2. Materials and Methods

### 2.1 Preparation of Advanced Glycation End-Products

AGE-modified bovine serum albumin (MG-BSA) was prepared following a modified protocol as previously described in Matou-Nasri et al. (2017). Briefly, BSA (Fraction V; Sigma-Aldrich) was incubated with 0.1 M methylglyoxal (Sigma-Aldrich) in 0.1 M sodium phosphate buffer (pH 7.4) containing 0.1% sodium azide at 37°C for 72 h. Control BSA was prepared identically without methylglyoxal. Both preparations were dialysed against distilled water and passed through Detoxi-Gel Endotoxin Removing Gel columns (Thermo Scientific); endotoxin levels were confirmed below detection threshold (<0.125 EU/mL) using an E-Toxate Limulus Amebocyte Lysate assay. AGE formation was verified spectrofluorometrically (excitation 350 nm, emission 420 nm; Perkin Elmer LS 30). Protein concentration was determined by Bradford assay (Bio-Rad).

### 2.2 Human Neuromuscular Junction Co-Culture Model and AGE Treatment

A fully human, serum-free NMJ co-culture system was established using GFP-labelled neural progenitor cells (NPCs), obtained in 2015 and previously described by Abd Al Samid et al. (2018), together with C83 immortalised human myoblasts obtained in 2014 from the MyoLine immortalisation platform of the Institut de Myologie (Paris, France). The C83 human immortalised myoblast cell line was derived from a semitendinosus muscle biopsy of a healthy 83-year-old male donor through Myobank, a tissue bank affiliated with EuroBioBank and authorised by the French Ministry of Research (authorisation AC-2013-1868). The cells were immortalised using hTERT and CDK4 as described by Mamchaoui et al. (2011). No Research Resource Identifier (RRID) was identified for either the C83 myoblast line or the GFP-labelled NPCs following searches of publicly available cell line databases. STR authentication data were not available for the C83 myoblast line. The C83 cell line has been previously characterised for its proliferative capacity and myogenic differentiation potential (Mamchaoui et al., 2011), has not been reported as misidentified or cross-contaminated in the International Cell Line Authentication Committee (ICLAC) Register or the Cellosaurus problematic cell line database, and has been subsequently applied in the human NMJ co-culture model(Abd Al Samid et al., 2018; Jasdeep Saini et al., 2020). Cells were routinely screened for mycoplasma contamination, and all experiments were performed using mycoplasma-free cultures. The C83 line was selected to provide an aged human myogenic component relevant to modelling neuromuscular ageing. Cells were co-cultured in serum-free differentiation medium for up to 12 days. Full culture details are provided in Supplementary Methods S1.

On day 3 of co-culture, cultures were treated with AGEs (200 µg/mL), non-modified BSA (200 µg/mL), or differentiation medium alone. The working concentration of 200 µg/mL was selected based on a dose-response experiment (50-200 µg/mL) showing the most consistent impairment in axonal morphology and mitochondrial bioenergetics at this dose. Treatments were maintained throughout the differentiation period, with medium refreshed every 48 hours until day 12.

### 2.3 Immunocytochemistry and Morphological Analysis

On day 12 of co-culture, cells were fixed in 4% paraformaldehyde (10 min at room temperature), permeabilised with 0.5% Triton X-100 in PBS (15 min), and blocked in PBS containing 1% BSA and 10% goat serum (1 h). Cells were incubated with anti-Myosin Heavy Chain (MF20, eFluor 660; eBioscience, AB_2574267; 1:400) for 1 h at room temperature. For model validation, cells were additionally incubated with anti-βIII-tubulin (eBioscience, AB_2574421; 1:400). Nuclei were counterstained with DAPI (1:10,000). For NMJ validation, live co-cultures were incubated with α-bungarotoxin-Alexa Fluor™ 647 (1 µg/mL; Invitrogen, B35450) for 30 min prior to fixation as described in Supplementary Methods S2. Images were acquired using a Leica DMI6000B inverted fluorescence microscope or Leica STELLARIS 5 confocal microscope. At least 10 random fields per condition were analysed. Myotube morphology was quantified from MyHC-stained images using ImageJ, including MyHC-positive area, area fraction, mean myotube area, and minimum Feret diameter. Fusion index was calculated as the percentage of nuclei within myotubes containing ≥3 nuclei. Axonal length was measured from GFP-positive axons using Simple Neurite Tracer plugin in ImageJ.

### 2.4 Mitochondrial Function Assessment

Mitochondrial membrane potential (ΔΨm), mitochondrial mass, and superoxide production were assessed on day 12 of co-culture using TMRM (200 nM; Thermo Fisher Scientific), MitoTracker™ Green FM (100 nM; Thermo Fisher Scientific), and MitoSOX™ Red (5 µM; Invitrogen), respectively, as previously described (Thoma et al., 2020). MitoSOX-stained cultures were imaged live; TMRM and MitoTracker-stained cultures were fixed in 4% paraformaldehyde prior to imaging. Fluorescence images were acquired using a Leica STELLARIS 5 Confocal Microscope (40× oil-immersion objective). At least 10 random fields per condition were imaged, and fluorescence intensity quantified using ImageJ (v1.53k). TMRM fluorescence was normalised to MitoTracker Green signal to account for differences in mitochondrial content.

### 2.5 Mitochondrial Bioenergetic Analysis (Seahorse XF Mito Stress Test)

Mitochondrial respiration was assessed using the Agilent Seahorse XFp Analyser (Agilent Technologies, CA, USA) as previously described (Thoma et al., 2025). Co-cultures were seeded into XFp Miniplates (7,000 cells/well) and treated with AGEs or BSA, for 9 days. On the day of the assay, the medium was replaced with non-buffered XF Assay Medium (10 mM glucose, 10 mM pyruvate, 2 mM glutamine, pH 7.4), and cells were equilibrated for 1 h at 37°C. Sequential injections of oligomycin (1 µM), FCCP (4 µM), and rotenone/antimycin A (1 µM each) were performed using the Seahorse XFp Cell Mito Stress Test Kit. OCR was normalised to total protein content by BCA assay and expressed as pmol O₂ min⁻¹ µg⁻¹ protein. Derived parameters included basal respiration, ATP-linked respiration, maximal respiration, spare respiratory capacity, proton leak, and non-mitochondrial respiration.

### 2.6 Quantitative Analysis of Myotube Contraction

Spontaneous myotube contraction intensity and frequency were quantified by video microscopy on day 12 of co-culture as described in Supplementary Methods S3. Representative contraction videos from BSA-treated and AGE-treated co-cultures are provided as Supplementary Videos S2 and S3.

### 2.7 Human Growth Factor Array

Conditioned media from AGE-treated, BSA-treated, and untreated co-cultures were collected on day 12, centrifuged to remove debris. Forty neurotrophic and myogenic growth factors were simultaneously quantified using a Quantibody® Human Growth Factor Array (RayBiotech, Norcross, GA, USA) according to the manufacturer’s instructions. Arrays were scanned using a GenePix 4000B laser scanner (Molecular Devices) at 532 nm, and raw fluorescence intensities extracted using GenePix Pro 4.1 software. Data were background-subtracted, normalised to internal positive controls, and expressed as relative fluorescence units (RFU).

### 2.8 Statistical Analysis

Statistical analyses were performed using GraphPad Prism v9.4.1. Data were tested for normality using the Shapiro–Wilk test and homogeneity of variance using Levene’s test. For two-group comparisons, an unpaired Student’s *t*-test was used. For multiple-group comparisons, one-way ANOVA with Tukey’s post hoc test was used. Data are presented as mean ± SD. Statistical significance was accepted at *p* < 0.05.

## 3. Results

### 3.1 AGEs Induce Reduced Axonal Length and Myotube Remodelling

AGE exposure induced significant structural changes in motor neurons and myotubes by day 12 (Figure 1). Myotube fusion index, mean myotube area, and mean myotube width were all significantly greater in AGE-treated cultures compared to BSA controls (Figures 1C-E). MyHC-positive area fraction did not differ significantly between conditions (Figure 1B). Motor neuron axonal length was significantly reduced in AGE-treated cultures compared to BSA controls (Figure 1F).

**Figure 1:**
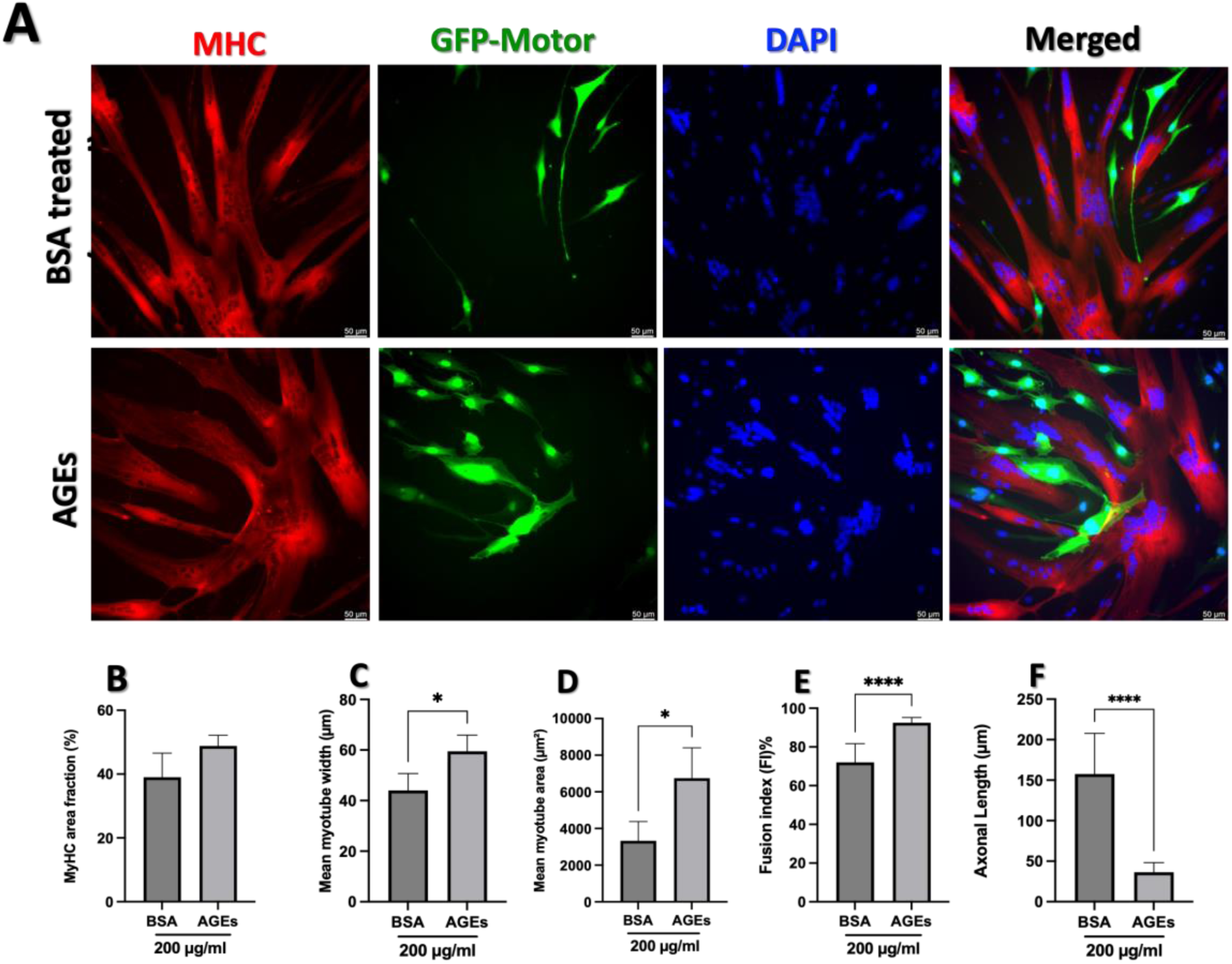
Effects of AGE exposure on neuronal and myotube morphology in the human NMJ co-culture. (A) Representative immunofluorescence images of BSA-treated (control) and AGE-treated co-cultures at day 12, stained for myosin heavy chain (MHC; red) and GFP-expressing motor neurons (green); nuclei counterstained with DAPI (blue). AGE treatment reduced axonal outgrowth and induced myotube enlargement with centralised nuclear clustering. Scale bar = 50 µm. Quantification of (B) MyHC-positive area fraction (%), (C) mean myotube width (µm), (D) mean myotube area (µm²), (E) fusion index (%), and (F) axonal length (µm). Data are mean ± SD; unpaired t-test; \**p* < 0.05, \*\*\*\**p* < 0.0001. n = 3 independent experiments.

### 2.2 AGEs Impair Mitochondrial Bioenergetics in the Human NMJ Co-culture

Basal and maximal respiration did not differ significantly between conditions (Figure 2A, C-D). Real-time ECAR measurements showed lower extracellular acidification in AGE-treated cultures than in BSA controls across all time points (Figure 2B). AGE treatment was associated with a significant increase in proton leak (*p* = 0.002; Figure 2E), a significant reduction in ATP-linked respiration (*p* = 0.0179; Figure 2F), and a significant reduction in spare respiratory capacity (*p* = 0.0017; Figure 2G).

**Figure 2:**
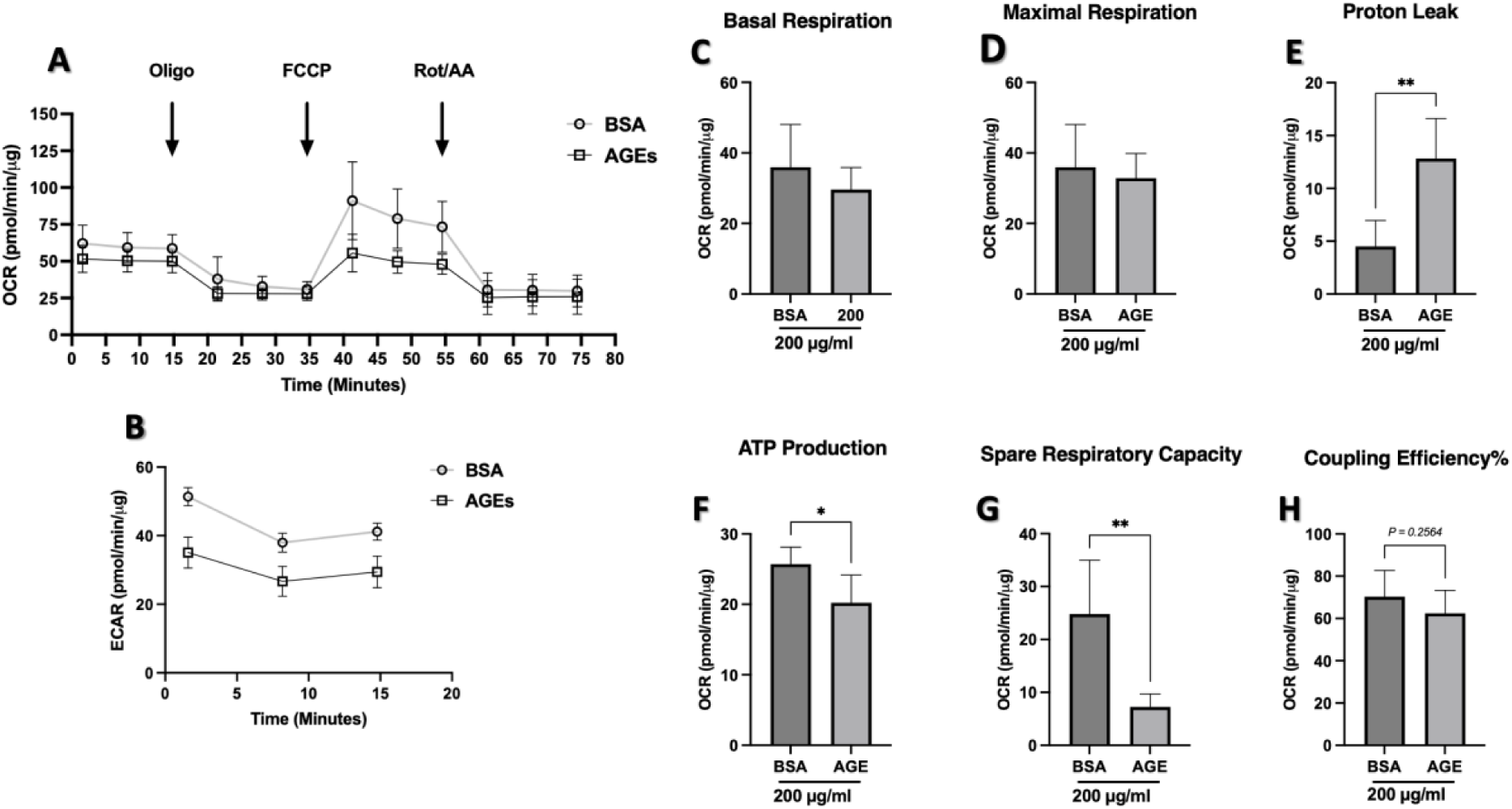
Effects of AGEs on mitochondrial bioenergetics in the human NMJ co-culture. (A) Real-time OCR following sequential injection of oligomycin, FCCP, and rotenone/antimycin A. (B) Real-time ECAR. Derived parameters: (C) basal respiration, (D) maximal respiration, (E) proton leak, (F) ATP-linked respiration, (G) spare respiratory capacity, and (H) coupling efficiency, normalised to protein content. Data are mean ± SD; unpaired t-test; \**p* < 0.05, \*\**p* < 0.01. n = 4 independent experiments.

### 2.3 AGEs Depolarise Mitochondrial Membrane Potential and Elevate Mitochondrial Superoxide

TMRM fluorescence was significantly reduced in AGE-treated cultures compared to BSA controls (*p* = 0.0103; Figure 3B), and TMRM/MitoTracker ratio was significantly reduced (*p* = 0.0015; Figure 3C), confirming mitochondrial membrane depolarisation independent of changes in mitochondrial mass. MitoSOX Red fluorescence was significantly elevated in AGE-treated cultures (*p* = 0.0008; Figure 3E), and MitoSOX/MitoTracker ratio was significantly elevated (*p* = 0.0001; Figure 3F), confirming increased superoxide production per unit mitochondrial mass. MitoTracker Green fluorescence did not differ significantly between conditions, indicating no change in mitochondrial mass.

**Figure 3:**
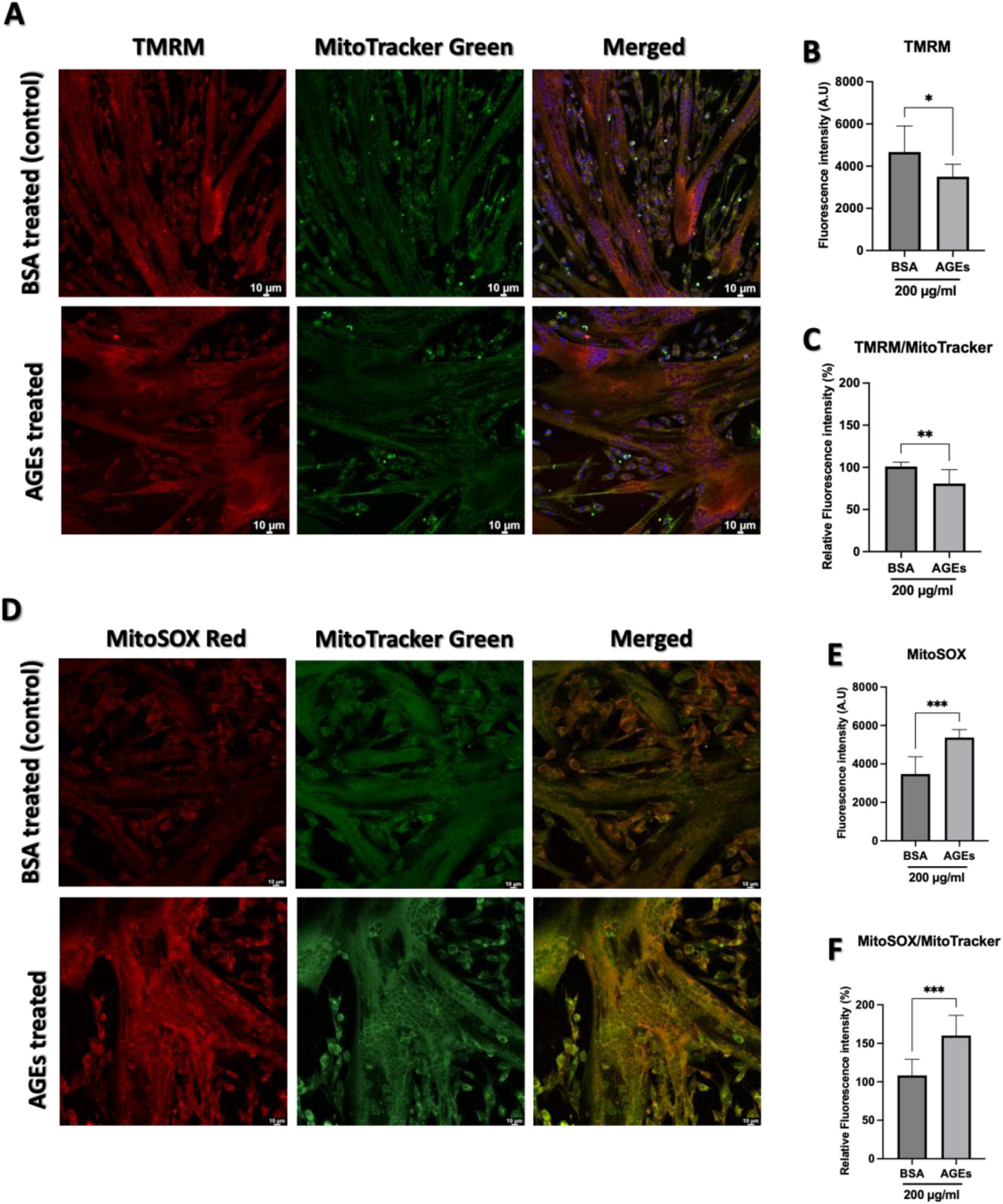
Effects of AGEs on mitochondrial membrane potential and superoxide production. (A) Representative fluorescence images of BSA-treated and AGE-treated co-cultures stained with TMRM (red) and MitoTracker Green (green). (B) TMRM fluorescence intensity and (C) TMRM/MitoTracker ratio. (D) Representative fluorescence images stained with MitoSOX Red and MitoTracker Green. (E) MitoSOX Red fluorescence intensity and (F) MitoSOX/MitoTracker ratio. Data are mean ± SD; unpaired t-test; \**p* < 0.05, \*\**p* < 0.01, \*\*\**p* < 0.001. n = 3 independent experiments. arbitrary unit (A.U.).

### 2.4 AGEs Reduce Spontaneous Contractile Activity in the Human NMJ Co-culture

Mean contraction intensity was significantly reduced in AGE-treated cultures compared to BSA controls (0.034 ± 0.009 a.u. vs 0.053 ± 0.01 a.u.; *p* = 0.0002; Figure 4C), as was spontaneous contraction frequency (10.19 ± 3 s⁻¹ vs 15.35 ± 2 s⁻¹; *p* = 0.0017; Figure 4D).

**Figure 4:**
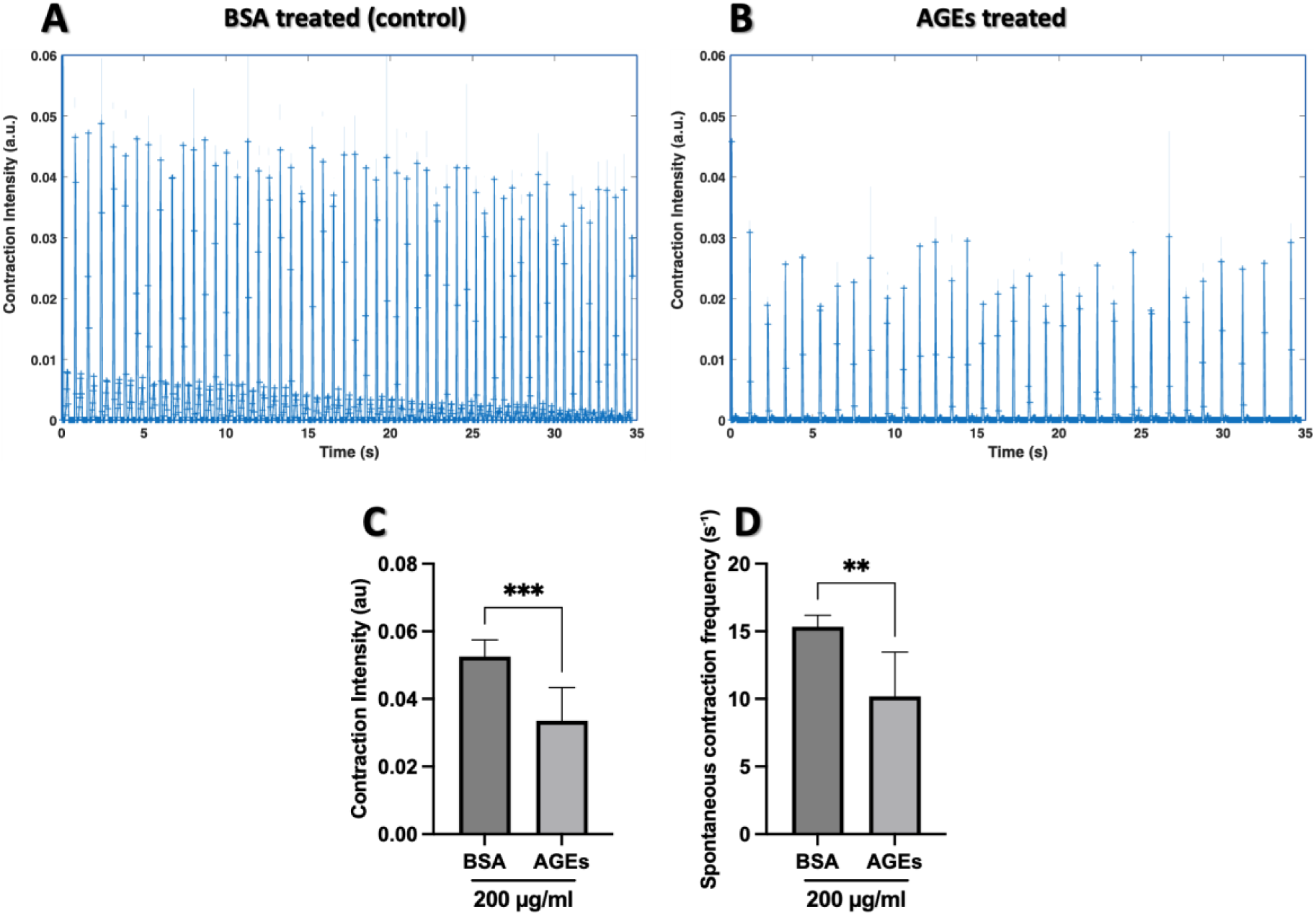
Effects of AGEs on spontaneous contractile activity. Representative motion-analysis traces of contraction intensity over time in (A) BSA-treated and (B) AGE-treated co-cultures on day 12, recorded over 35 seconds. Quantification of (C) mean contraction intensity (a.u.) and (D) spontaneous contraction frequency (s⁻¹). Data are mean ± SD; unpaired t-test; \*\**p* < 0.01, \*\*\**p* < 0.001. n = 5 independent experiments. s⁻¹: contractions per second.

### 2.5 AGEs Suppress Neurotrophic and Myogenic Growth Factor Secretion in the Human NMJ Co-culture

Growth factor array analysis of conditioned media revealed significant reductions in eight detected factors in AGE-treated cultures compared to BSA controls, including BDNF (*p* = 0.0435; Figure 5A), GDF-15 (*p* = 0.0320; Figure 5B), IGFBP-3 (*p* = 0.0277; Figure 5C), NGFR (*p* = 0.0071; Figure 5D), bFGF (*p* = 0.0203; Figure 5E), GDNF (*p* = 0.0013; Figure 5F), IGF (*p* = 0.0148; Figure 5G), and NT3 (*p* = 0.0003; Figure 5H).

**Figure 5:**
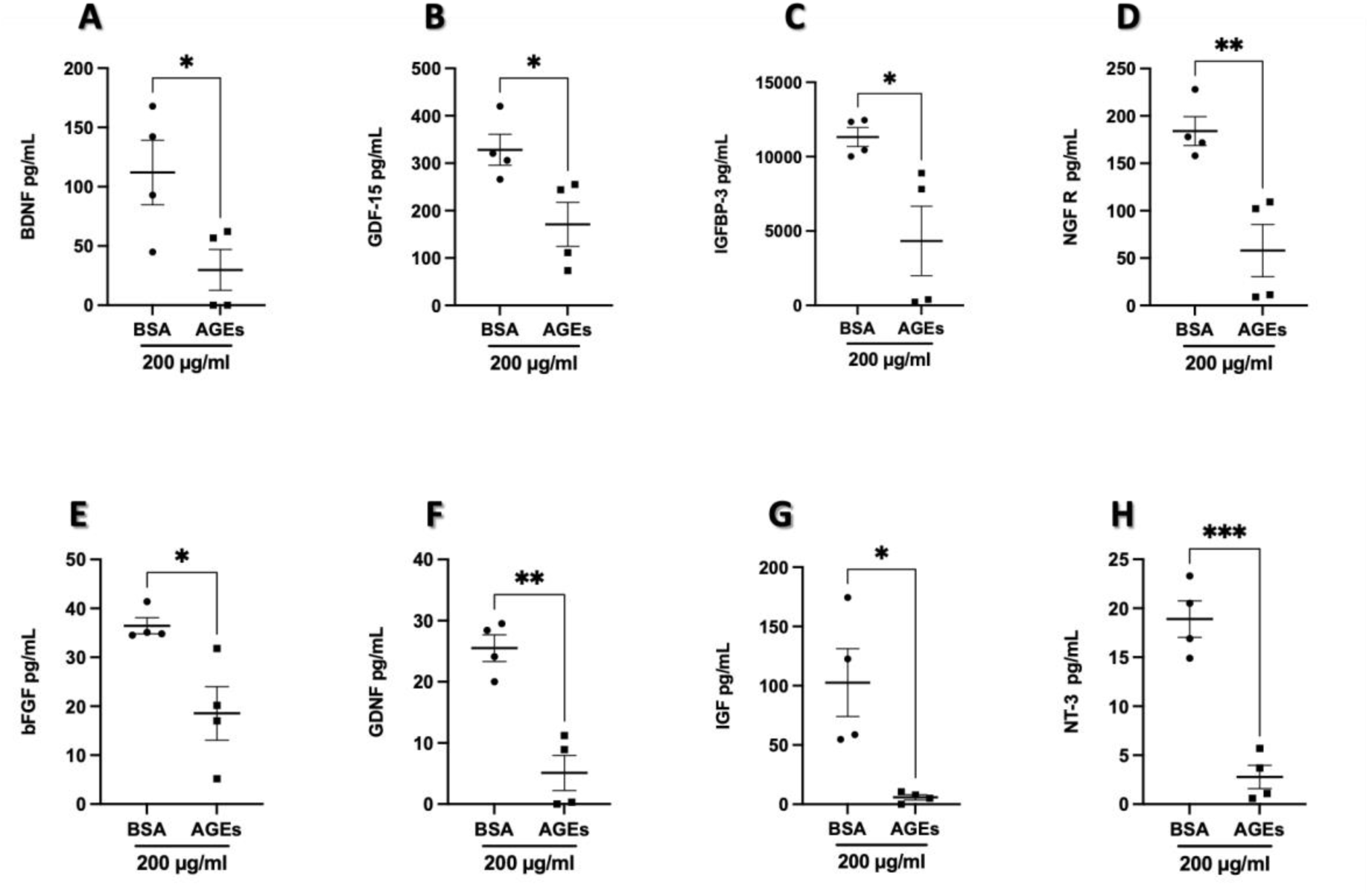
Effects of AGEs on neurotrophic and myogenic factor secretion. Concentrations of (A) BDNF, (B) GDF-15, (C) IGFBP-3, (D) NGFR, (E) bFGF, (F) GDNF, (G) IGF, and (H) NT3 (all pg/mL) in conditioned media from BSA-treated and AGE-treated co-cultures at day 12, determined by growth factor antibody array. Data are mean ± SD; unpaired t-test; \**p* < 0.05, \*\**p* < 0.01, \*\*\**p* < 0.001. n = 3 independent experiments.

## 3. Discussion

This study employed a fully human, serum-free and neural growth factor-free NMJ co-culture system to investigate the effects of AGE exposure on neuromuscular integrity. AGE treatment induced axonal impairment, myotube remodelling, mitochondrial uncoupling, reduced spontaneous contractile activity, and suppression of neurotrophic and myogenic growth factor secretion. Together, these findings suggest that the NMJ microenvironment is sensitive to glycation stress and that AGE exposure disrupts several interconnected features required for neuromuscular function.

Importantly, these findings are relevant to conditions in which AGE accumulation is increased, including ageing, type 2 diabetes, chronic kidney disease, and sarcopenia. In these settings, muscle weakness is unlikely to arise from myofibre pathology alone but rather reflects a combined disruption of muscle metabolism, motor neuron input, extracellular matrix integrity, and trophic signalling. The present data support this concept by showing that AGE exposure affects multiple components of the NMJ microenvironment in parallel. This positions glycation stress as a potential contributor to neuromuscular decline rather than solely a modifier of skeletal muscle structure.

A prominent structural finding was the marked reduction in motor neuron axonal length following AGE exposure. Mechanistically, this may be associated with glycation-mediated crosslinking of neurofilament subunits and tubulin into insoluble aggregates, as reported in methylglyoxal-exposed neuronal cells, directly disrupting cytoskeletal integrity and impairing axonal transport (Ooi, Furukawa, Takeuchi, & Koriyama, 2024). AGE modification of extracellular matrix proteins, including laminin and fibronectin, has also been shown to impair integrin-mediated adhesion and reduce neurotrophin-stimulated neurite outgrowth, suggesting that glycation stress can affect both neuronal structure and the matrix environment required for axonal extension (Bennmann et al., 2014; Duran-Jimenez et al., 2009). AGE-RAGE signalling may further contribute by activating the NF-κB and MAPK pathways, which are known to alter inflammatory and stress-responsive gene expression (Ott et al., 2014). Since motor neurons and myotubes share the same medium in the present model, it is not possible to determine whether reduced axonal length reflects direct AGE toxicity to the neuronal population or an indirect consequence of altered myotube-derived trophic support. However, reduced axonal extension would be expected to decrease synaptic contact with myotubes, consistent with the observed reductions in spontaneous contraction intensity and frequency.

AGE exposure also produced enlarged myotubes with increased fusion index, area, and width, without a corresponding increase in MyHC-positive area fraction. This indicates that AGE-treated cultures contained larger and more fused myotubes, but not a greater total MyHC-positive area. This pattern does not necessarily indicate physiological hypertrophy, which would require increased accumulation of contractile proteins and improved functional output. Instead, the persistence of centralised nuclei suggests impaired terminal maturation. During normal myotube formation, nuclear positioning depends on microtubule organisation and motor protein activity, and disruption of this process is linked to defective myofibre maturation and weakness (Folker & Baylies, 2013; Gache, Gomes, & Cadot, 2017). Notably, age-related deterioration of the microtubule network has been directly linked to disordered nuclear positioning in aged muscle, suggesting that AGE accumulation may amplify an age-intrinsic vulnerability of the nuclear positioning apparatus (Cisterna & Malatesta, 2024). Centralised nuclei are not merely a marker of failed maturation; mispositioned nuclei disrupt myonuclear domain organisation and local gene product delivery and have been identified as a direct contributor to muscle weakness (Folker & Baylies, 2013). Thus, the structural phenotype observed here suggests that AGEs impair both neuronal outgrowth and myotube maturation, providing a morphological basis for the functional decline observed in the co-culture.

Bioenergetic analysis showed that AGE exposure reduced ATP-linked respiration and spare respiratory capacity while increasing proton leak, without significant changes in basal or maximal respiration. This profile is consistent with mitochondrial uncoupling rather than a generalised collapse of respiratory function. In this context, oxygen consumption is maintained, but a greater proportion of respiration is diverted away from ATP production due to increased proton leakage across the inner mitochondrial membrane (Divakaruni & Jastroch, 2022). Similar AGE-induced metabolic disruption has been reported in skeletal muscle cells exposed to methylglyoxal, in which mitochondrial respiration and ATP production were reduced, accompanied by increased oxidative stress (Du et al., 2023; Todoriki et al., 2022). In the present study, ECAR did not increase to compensate for reduced ATP-linked respiration, suggesting that glycolytic activity did not offset the mitochondrial deficit. This supports the interpretation that AGE exposure compromises bioenergetic flexibility in the NMJ co-culture. The reduction in spare respiratory capacity is particularly relevant for neuromuscular function, as both motor neurons and myotubes require rapid bioenergetic adaptation during synaptic transmission and contraction. Mitochondria at the NMJ contribute to ATP supply and calcium buffering, processes that are essential for synaptic vesicle cycling and excitation-contraction coupling (Davis et al., 2022). However, respiration measurements in the present study reflect the combined metabolic status of the entire co-culture population and cannot distinguish neuronal from myogenic mitochondrial responses. Recent work shows that postsynaptic mitochondrial disruption contributes to NMJ fragmentation and impaired reinnervation (Guzman et al., 2026), though the relative contributions of each cell type in the present model remain unresolved.

Consistent with the Seahorse data, fluorescence-based analysis showed reduced TMRM/MitoTracker ratio and increased MitoSOX/MitoTracker ratio, indicating mitochondrial depolarisation and increased mitochondrial superoxide production per unit mitochondrial mass. Proton leak can promote electron leakage from complexes I and III and increase superoxide generation rather than ATP synthesis, consistent with the co-occurrence of mitochondrial uncoupling and elevated mitochondrial ROS observed here (Murphy, 2009).

The functional consequences of AGE exposure were evident in the significant reductions in spontaneous contraction intensity and frequency. In this model, spontaneous contractions depend on functional communication between motor neurons and myotubes, as supported by validation experiments showing inhibition by α-bungarotoxin and potentiation by L-glutamic acid (Jasdeep Saini et al., 2020); Supplementary Figure S2). The reduced contractile output, therefore, aligns with the structural and metabolic alterations observed in AGE-treated co-cultures. While AChR clustering was not assessed under AGE treatment conditions, the combination of shortened axons, mitochondrial dysfunction, and reduced contraction supports impaired neuromuscular function. These findings also align with clinical and experimental evidence linking AGE accumulation to reduced muscle strength, impaired physical function, and sarcopenia, including in older adults and individuals with type 2 diabetes (Granic et al., 2023; Mori et al., 2019). The magnitude of functional impairment observed here is comparable to that reported in AGE-treated 3D skeletal muscle models, extending previous muscle-only findings to a human NMJ co-culture system (Kim et al., 2025).

AGE treatment also significantly reduced levels of several detectable neurotrophic and myogenic growth factors measured in conditioned media, including BDNF, GDNF, NT3, NGFR, bFGF, IGF, and IGFBP-3. Cell viability was confirmed across all AGE treatment conditions prior to conditioned media collection, indicating that the observed reductions reflect impaired biosynthesis or secretory activity rather than loss of cell number. Since the culture system is serum-free and neural growth factor-free, these signals reflect endogenous secretion from the co-culture. BDNF, GDNF, and NT3 are important regulators of motor neuron survival, axonal growth, and NMJ maintenance (Gonzalez-Freire et al., 2014). Their suppression raises the possibility that reduced axonal length may reflect trophic withdrawal rather than, or in addition to, direct glycation-related neuronal stress. The reduction in NGFR is also notable, as NGFR is involved in mature NMJ organisation, synaptic vesicle availability, postsynaptic AChR organisation, and force production (Pérez et al., 2019). Similarly, reduced IGF, IGFBP-3, and bFGF may contribute to impaired myotube maturation by limiting anabolic and myogenic signalling (Baoge et al., 2012; Ilyushchenko, Matchekhina, Melnitskaia, & Strazhesko, 2024). The uniform suppression of both neural and myogenic factors suggests a broad reduction in secretory output rather than selective disruption of a single pathway. AGE-mediated protein modification or RAGE-dependent transcriptional changes could each contribute to this phenotype, but the present study did not directly isolate these mechanisms.

One consideration of the present model is that motor neurons and myotubes share the same co-culture environment; therefore, the relative contribution of each cell type cannot be separated. AGE exposure also represents an acute glycation challenge rather than lifelong tissue accumulation. Nevertheless, this design preserves reciprocal neuromuscular communication in a fully human, serum-free system without exogenous growth or neurotrophic factors. Overall, AGEs exert multi-level detrimental effects on the integrity of human NMJ co-cultures, including axonal impairment, abnormal myotube maturation, mitochondrial uncoupling, reduced contractile activity, and suppression of trophic signalling. These findings support the use of this platform to investigate glycation-related neuromuscular decline and test interventions targeting AGE accumulation, RAGE signalling, mitochondrial stress, or trophic support.

## Author Contributions

*Conception and experimental design*: Razan Alomosh, Nasser Al-Shanti, Adam P. Lightfoot, Nessar Ahmed. *Data collection*: Razan Alomosh, Amy Bateman. *Data analysis and interpretation*: Razan Alomosh, Amy Bateman, Nasser Al-Shanti, Adam P. Lightfoot, Nessar Ahmed. *Ressources*: Kamel Mamchaoui, Vincent Mouly, Moi Hoon Yap*. Software and Methodology:* Moi Hoon Yap. *Manuscript writing*: Razan Alomosh, Adam P. Lightfoot, Nasser Al-Shanti. *Final approval of manuscript*: All authors.

## Conflict of interest statement

The authors declare no conflict of interest.

## Acknowledgements

The authors thank the Department of Life Sciences and the Department of Computing and Mathematics, Faculty of Science and Engineering, Manchester Metropolitan University (Manchester, UK) for providing the facilities and resources that supported this research. The authors also thank the MyoLine platform of the Institut de Myologie (Paris, France) for providing the C83 immortalised human myoblast cell line.

## Data availability statement

All data generated or analysed during this study are included in this published article. The custom MATLAB code (version R2022a, 9.12.0) used for quantitative analysis of myotube contraction is available from the corresponding author upon reasonable request.

## Notes

### Competing Interest Statement

The authors have declared no competing interest.

## References

Abd Al Samid, M., McPhee, J. S., Saini, J., McKay, T. R., Fitzpatrick, L. M., Mamchaoui, K.,…Al-Shanti, N. (2018). A functional human motor unit platform engineered from human embryonic stem cells and immortalized skeletal myoblasts. Stem Cells Cloning, 11, 85–93. doi:10.2147/sccaa.S178562

Adachi, N., Kanazawa, I., Tanaka, K.-i., Takeno, A., Notsu, M., Tanaka, S., & Sugimoto, T. (2019). Insulin-Like Growth Factor-I Protects Against the Detrimental Effects of Advanced Glycation End Products and High Glucose in Myoblastic C2C12 Cells. Calcified Tissue International, 105(1), 89–96. doi:10.1007/s00223-019-00537-w

Baoge, L., Van Den Steen, E., Rimbaut, S., Philips, N., Witvrouw, E., Almqvist, K. F.,…Vanden Bossche, L. C. (2012). Treatment of skeletal muscle injury: a review. ISRN Orthop, 2012, 689012. doi:10.5402/2012/689012

Bennmann, D., Horstkorte, R., Hofmann, B., Jacobs, K., Navarrete-Santos, A., Simm, A.,…Gnanapragassam, V. S. (2014). Advanced glycation endproducts interfere with adhesion and neurite outgrowth. PLoS One, 9(11), e112115. doi:10.1371/journal.pone.0112115

Chai, S., Zhang, N., Cui, C., Bao, Z., Wang, Q., Lin, W.,…Cheung, W. H. (2026). Systematic review of mitochondrial dysfunction and oxidative stress in aging: A focus on neuromuscular junctions. Neural Regeneration Research, 21(5), 1947–1960. doi:10.4103/nrr.Nrr-d-24-01338

Chiu, C. Y., Yang, R. S., Sheu, M. L., Chan, D. C., Yang, T. H., Tsai, K. S.,…Liu, S. H. (2016). Advanced glycation end-products induce skeletal muscle atrophy and dysfunction in diabetic mice via a RAGE-mediated, AMPK-down-regulated, Akt pathway. The Journal of pathology, 238(3), 470–482.

Cisterna, B., & Malatesta, M. (2024). Molecular and Structural Alterations of Skeletal Muscle Tissue Nuclei during Aging. Int J Mol Sci, 25(3). doi:10.3390/ijms25031833

Davis, L. A., Fogarty, M. J., Brown, A., & Sieck, G. C. (2022). Structure and Function of the Mammalian Neuromuscular Junction. Compr Physiol, 12(4), 3731–3766. doi:10.1002/cphy.c210022

Divakaruni, A. S., & Jastroch, M. (2022). A practical guide for the analysis, standardization and interpretation of oxygen consumption measurements. Nat Metab, 4(8), 978–994. doi:10.1038/s42255-022-00619-4

Du, H., Ma, Y., Wang, X., Zhang, Y., Zhu, L., Shi, S.,…Liu, Z. (2023). Advanced glycation end products induce skeletal muscle atrophy and insulin resistance via activating ROS-mediated ER stress PERK/FOXO1 signaling. Am J Physiol Endocrinol Metab, 324(3), E279–e287. doi:10.1152/ajpendo.00218.2022

Duran-Jimenez, B., Dobler, D., Moffatt, S., Rabbani, N., Streuli, C. H., Thornalley, P. J.,…Gardiner, N. J. (2009). Advanced glycation end products in extracellular matrix proteins contribute to the failure of sensory nerve regeneration in diabetes. Diabetes, 58(12), 2893–2903. doi:10.2337/db09-0320

Folker, E., & Baylies, M. (2013). Nuclear positioning in muscle development and disease. Frontiers in physiology, Volume 4 - 2013. doi:10.3389/fphys.2013.00363

Gache, V., Gomes, E. R., & Cadot, B. (2017). Microtubule motors involved in nuclear movement during skeletal muscle differentiation. Mol Biol Cell, 28(7), 865–874. doi:10.1091/mbc.E16-06-0405

Gonzalez-Freire, M., de Cabo, R., Studenski, S. A., & Ferrucci, L. (2014). The Neuromuscular Junction: Aging at the Crossroad between Nerves and Muscle. Frontiers in aging neuroscience, Volume 6 - 2014. doi:10.3389/fnagi.2014.00208

Granic, A., Hurst, C., Dismore, L., Dodds, R. M., Witham, M. D., Robinson, S. M., & Sayer, A. A. (2023). Advanced glycation end products in skeletal muscle health and sarcopenia: A systematic review of observational studies. Mech Ageing Dev, 209, 111744. doi:10.1016/j.mad.2022.111744

Guo, Z., Li, H., Jiang, S., Rahmati, M., Su, J., Yang, S.,…Deng, Z. (2025). The role of AGEs in muscle ageing and sarcopenia. Bone Joint Res, 14(3), 185–198. doi:10.1302/2046-3758.143.Bjr-2024-0252.R1

Guzman, S. D., Fraczek, P. M., Itsani, K., Furati, E. K., Juros, D., Kenney, G.,…Aguilar, C. A. (2026). Age-Associated Dysregulation of Postsynaptic Mitochondria Perturbs Reinnervation Kinetics. Aging Cell, 25(1), e70355. doi:10.1111/acel.70355

Ilyushchenko, A. K., Matchekhina, L. V., Melnitskaia, A. A., & Strazhesko, I. D. (2024). Associations of IGF-1 and IGFBP-3 with Aging and the Development of Age-Associated Diseases. Advances in Gerontology, 14(3), 118–126. doi:10.1134/S2079057025600041

Kim, J., Kim, I. U., Lee, Z. F., Han, J., Ahn, J., Jo, Y.,…Jeon, J. S. (2025). Detrimental effects of advanced glycation end-products (AGEs) on a 3D skeletal muscle model in microphysiological system. Biosensors and Bioelectronics, 278, 117316. 10.1016/j.bios.2025.117316

Li, L., Xiong, W.-C., & Mei, L. (2018). Neuromuscular junction formation, aging, and disorders. Annual review of physiology, 80, 159–188.

Mamchaoui, K., Trollet, C., Bigot, A., Negroni, E., Chaouch, S., Wolff, A.,…Mouly, V. (2011). Immortalized pathological human myoblasts: towards a universal tool for the study of neuromuscular disorders. Skelet Muscle, 1, 34. doi:10.1186/2044-5040-1-34

Matou-Nasri, S., Sharaf, H., Wang, Q., Almobadel, N., Rabhan, Z., Al-Eidi, H.,…Ahmed, N. (2017). Biological impact of advanced glycation endproducts on estrogen receptor-positive MCF-7 breast cancer cells. Biochimica et Biophysica Acta (BBA) - Molecular Basis of Disease, 1863(11), 2808–2820. 10.1016/j.bbadis.2017.07.011

Miao, Y., Xie, L., Song, J., Cai, X., Yang, J., Ma, X.,…Xie, P. (2024). Unraveling the causes of sarcopenia: Roles of neuromuscular junction impairment and mitochondrial dysfunction. Physiol Rep, 12(1), e15917. doi:10.14814/phy2.15917

Mori, H., Kuroda, A., Ishizu, M., Ohishi, M., Takashi, Y., Otsuka, Y.,…Matsuhisa, M. (2019). Association of accumulated advanced glycation end-products with a high prevalence of sarcopenia and dynapenia in patients with type 2 diabetes. J Diabetes Investig, 10(5), 1332–1340. doi:10.1111/jdi.13014

Murphy, M. P. (2009). How mitochondria produce reactive oxygen species. Biochem J, 417(1), 1–13. doi:10.1042/bj20081386

Olson, L. C., Redden, J. T., Schwartz, Z., Cohen, D. J., & McClure, M. J. (2021). Advanced Glycation End-Products in Skeletal Muscle Aging. Bioengineering, 8(11), 168. Retrieved from https://www.mdpi.com/2306-5354/8/11/168

Ooi, H., Furukawa, A., Takeuchi, M., & Koriyama, Y. (2024). Toxic Advanced Glycation End-Products Inhibit Axonal Elongation Mediated by β-Tubulin Aggregation in Mice Optic Nerves. Int J Mol Sci, 25(13). doi:10.3390/ijms25137409

Ott, C., Jacobs, K., Haucke, E., Navarrete Santos, A., Grune, T., & Simm, A. (2014). Role of advanced glycation end products in cellular signaling. Redox biology, 2, 411–429. 10.1016/j.redox.2013.12.016

Pérez, V., Bermedo-Garcia, F., Zelada, D., Court, F. A., Pérez, M., Fuenzalida, M.,…Henríquez, J. P. (2019). The p75(NTR) neurotrophin receptor is required to organize the mature neuromuscular synapse by regulating synaptic vesicle availability. Acta Neuropathol Commun, 7(1), 147. doi:10.1186/s40478-019-0802-7

Pratt, J., De Vito, G., Narici, M., & Boreham, C. (2020). Neuromuscular Junction Aging: A Role for Biomarkers and Exercise. The Journals of Gerontology: Series A, 76(4), 576–585. doi:10.1093/gerona/glaa207

Saini, J., Faroni, A., Reid, A. J., Mamchaoui, K., Mouly, V., Butler-Browne, G.,…Al-Shanti, N. (2020). A Novel Bioengineered Functional Motor Unit Platform to Study Neuromuscular Interaction. Journal of Clinical Medicine, 9(10), 3238. Retrieved from https://www.mdpi.com/2077-0383/9/10/3238

Saini, J., Faroni, A., Reid, A. J., Mouly, V., Butler-Browne, G., Lightfoot, A. P.,…Al-Shanti, N. (2021). Cross-talk between motor neurons and myotubes via endogenously secreted neural and muscular growth factors. Physiol Rep, 9(8), e14791. doi:10.14814/phy2.14791

Thoma, A., Alomosh, R., Bond, H. L., Akter-Miah, T., Al-Shanti, N., Degens, H.,…Lightfoot, A. P. (2025). A combination of major histocompatibility complex (MHC) I overexpression and type I interferon induce mitochondrial dysfunction in human skeletal myoblasts. Journal of cellular physiology, 240(1), e31458.

Thoma, A., Lyon, M., Al-Shanti, N., Nye, G. A., Cooper, R. G., & Lightfoot, A. P. (2020). Eukarion-134 Attenuates Endoplasmic Reticulum Stress-Induced Mitochondrial Dysfunction in Human Skeletal Muscle Cells. Antioxidants, 9(8), 710. Retrieved from https://www.mdpi.com/2076-3921/9/8/710

Todoriki, S., Hosoda, Y., Yamamoto, T., Watanabe, M., Sekimoto, A., Sato, H.,…Sato, E. (2022). Methylglyoxal Induces Inflammation, Metabolic Modulation and Oxidative Stress in Myoblast Cells. Toxins (Basel), 14(4). doi:10.3390/toxins14040263

Yipeng, X., Guiqian, W., Qiaochu, Z., Tengjie, H., Yan, Z., Hai, H., & Jing, Z. (2025). Molecular mechanisms by which mitochondrial dysfunction drives neuromuscular junction degeneration in amyotrophic lateral sclerosis. Neurobiology of disease, 216, 107103. 10.1016/j.nbd.2025.107103

Zgutka, K., Tkacz, M., Tomasiak, P., & Tarnowski, M. (2023). A Role for Advanced Glycation End Products in Molecular Ageing. Int J Mol Sci, 24(12). doi:10.3390/ijms24129881

Zhao, H., Iyama, R., Kurogi, E., Hayashi, T., & Egawa, T. (2024). Direct and acute effects of advanced glycation end products on proteostasis in isolated mouse skeletal muscle. Physiol Rep, 12(12), e16121. doi:10.14814/phy2.16121

