## Supplementary Information for "Detrimental effects of advanced glycation end-products (AGEs) on a human neuromuscular junction co-culture model"

Supplementary Figures S1- S2 and Supplementary Videos S1- S3 accompany this manuscript.

**Supplementary Figure S1. Structural validation of human neuromuscular junction co-culture model**

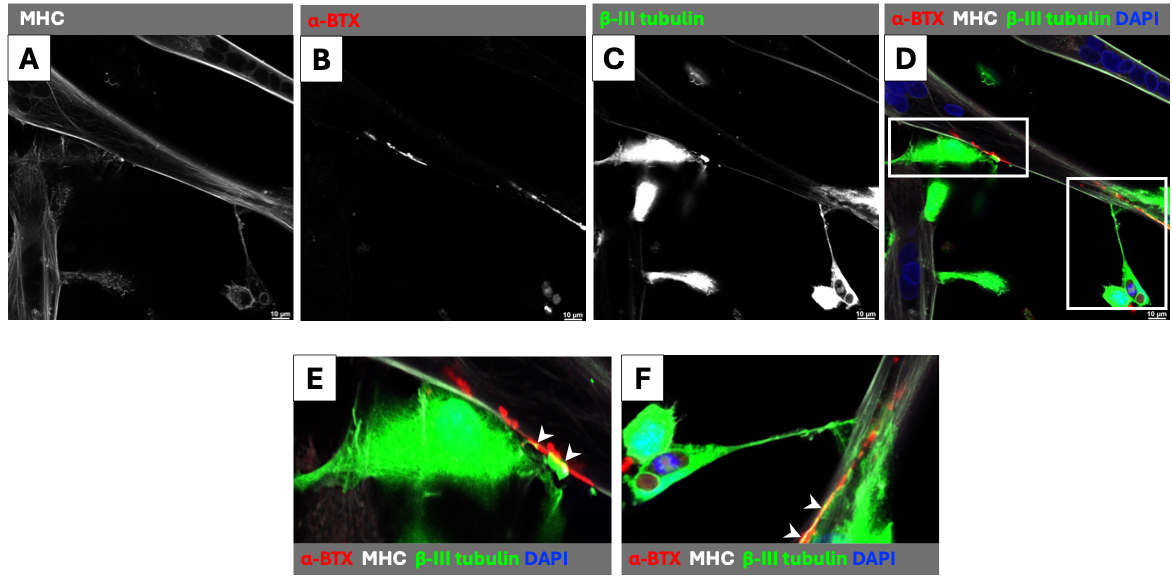

Spontaneous myotube contractions were confirmed by live video microscopy prior to fixation and staining. **(A)** Myosin heavy chain (MHC) immunostaining confirms myotube formation and morphology. **(B)** α-bungarotoxin (α-BTX) labelling of acetylcholine receptor (AChR) clusters at sites of motor neuron-myotube contact. **(C)** β-III tubulin immunostaining confirming motor neuron identity within the co-culture. **(D)** Merged overlay showing co-localisation of α-BTX-labelled AChR clusters (red), MHC-positive myotubes (grey), β-III tubulin-positive motor neurons (green), and DAPI nuclear counterstain (blue), confirming structural NMJ formation. **(E-F)** Magnified views of the boxed region in (D) illustrating co-localisation of AChR clusters with β-III tubulin-labelled nerve axons at sites of neuromuscular contact (indicated by white arrows). Scale bar = 10 µm.

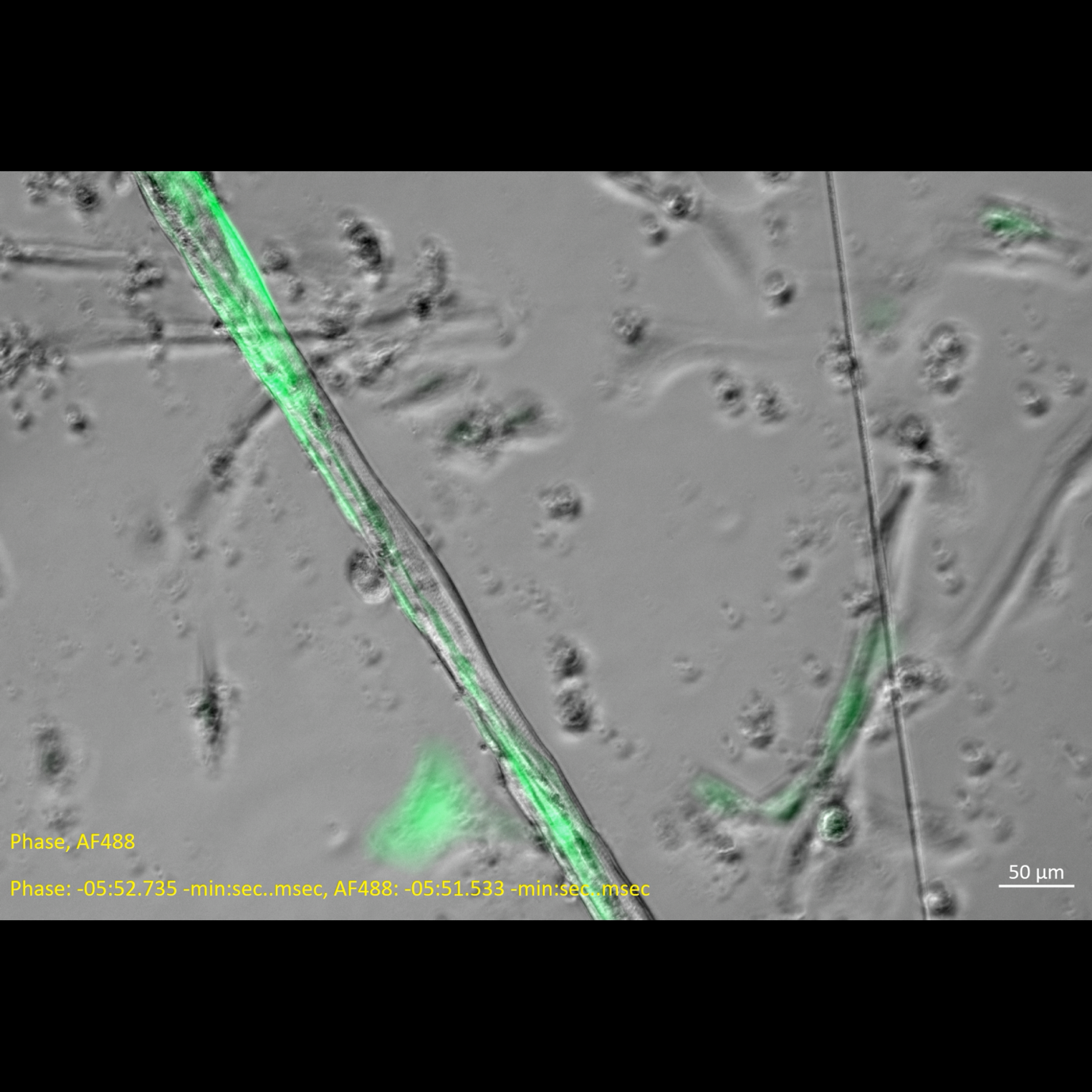
**Supplementary Video S1. Live time-lapse imaging of a GFP-labelled motor neuron in contact with a contracting myotube**

Overlay of GFP fluorescence and phase-contrast time-lapse video demonstrating live contractile activity in the NMJ co-culture. GFP-positive motor neuron axons can be seen in direct contact with spontaneously contracting myotubes, confirming functional neuromuscular connectivity. Scale bar = 50 µm. Video link: [Video.S1](https://mmutube.mmu.ac.uk/media/Supplementary+Video+S1.+Live+time-lapse+imaging+of+a+GFP-labelled+motor+neuron+in+contact+with+a+contracting+myotube/1_gbesraln)

**Supplementary Figure S2. Functional validation of human neuromuscular junction co-culture model**

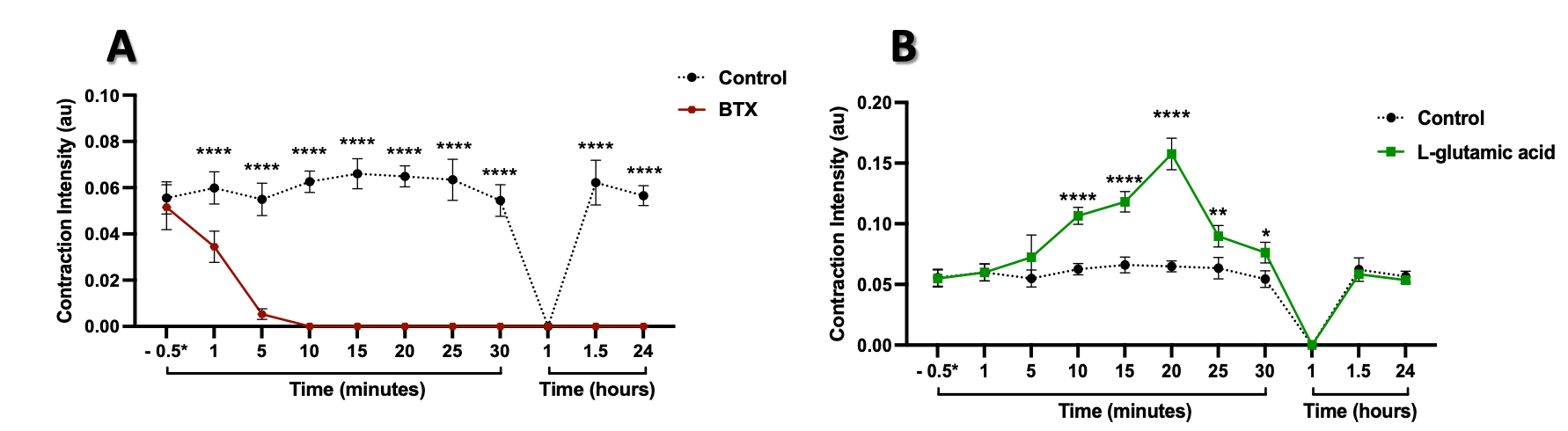

**(A)** Contraction intensity (CI) of myotubes monitored following application of α-bungarotoxin (α-BTX; 1 µg/mL, Alexa Fluor™ 647 conjugate, Invitrogen B35450) and compared with untreated controls over timepoints ranging from 30 seconds to 24 hours. α-BTX treatment resulted in rapid and complete abolition of contractile activity, with no recovery observed at any subsequent time point, including after medium replacement at 60 minutes and 24 hours, confirming that spontaneous contractions are dependent on acetylcholine receptor signalling at the NMJ. **(B)** Contraction intensity monitored following application of L-glutamic acid (400 µM; Sigma-Aldrich, G1251) compared with untreated controls over the same time points. L-glutamic acid produced a progressive increase in contraction intensity, peaking at 20 minutes and returning toward baseline by 30 minutes, with full recovery confirmed following medium replacement. Data represent three independent experiments (n = 3); CI is expressed in arbitrary units (au). Statistical comparisons at each time point were performed using two-way ANOVA with Šíčák’s multiple comparisons test. **p ≤ 0.01, ****p ≤ 0.0001 relative to control.

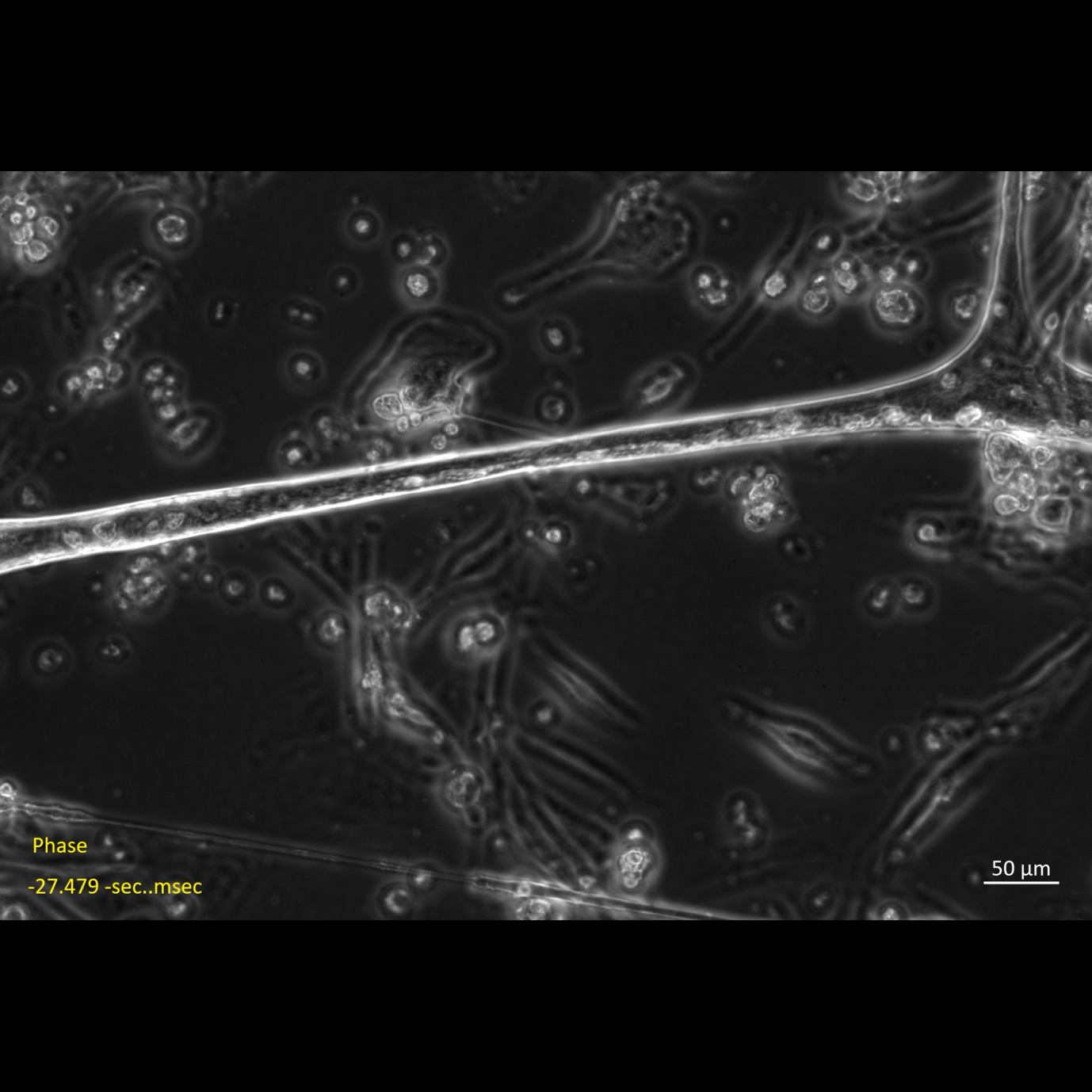
**Supplementary Video S2. Spontaneous myotube contraction in BSA-treated (control) co-culture**

Representative phase-contrast video of spontaneous myotube contractile activity in a BSA-treated co-culture (200 µg/mL) on day 12. Scale bar = 50 µm. Video link: [Video.S2](https://mmutube.mmu.ac.uk/media/Supplementary+Video+S2.+Spontaneous+myotube+contraction+in+BSA-treated+%28control%29+co-culture/1_etlbejwz)

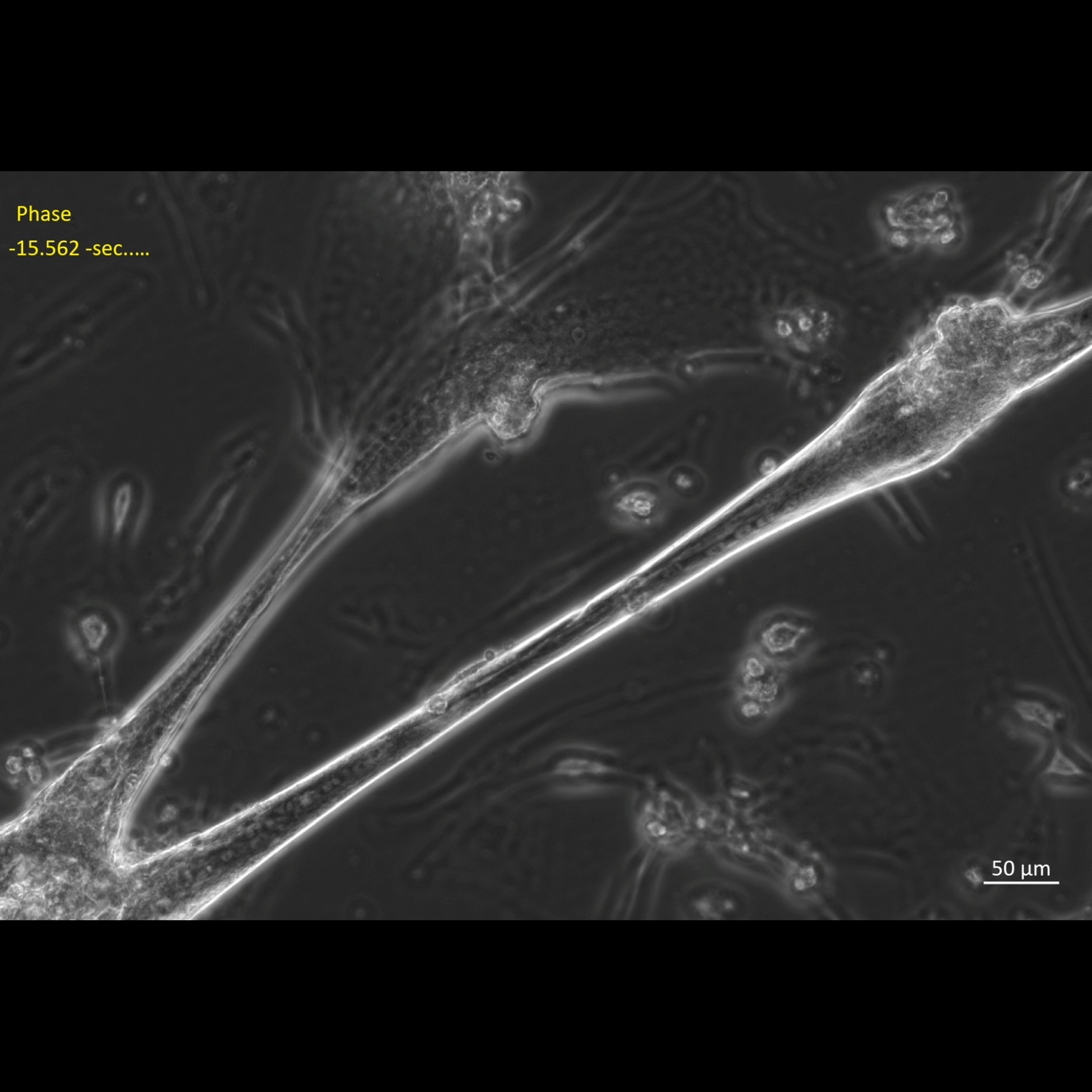
**Supplementary Video S3. Spontaneous myotube contraction in AGE-treated co-culture**

Representative phase-contrast video of spontaneous myotube contractile activity in an AGE-treated co-culture (200 µg/mL) on day 12, demonstrating visibly reduced contraction intensity and frequency compared to the BSA control. Scale bar = 50 µm. Video link: [Video.S3](https://mmutube.mmu.ac.uk/media/Supplementary+Video+S3.+Spontaneous+myotube+contraction+in+AGE-treated+co-culture/1_t4rlzw6n)

**Supplementary Methods S1. Cell Culture and NMJ Co-Culture Establishment**

GFP-labelled neural progenitor cells (NPCs), previously generated from the shef3 human embryonic stem cell line as described by Abd Al Samid et al. (2018), were used to enable identification and live tracking of motor neuron morphology. Stable GFP expression was confirmed prior to co-culture experiments. C83 myoblasts were cultured in growth medium composed of Medium 199 (HyClone, Fisher Scientific) and DMEM high-glucose (Corning, 15-013-CV) at a 1:4 ratio, supplemented with 20% heat-inactivated foetal bovine serum (Thermo Fisher Scientific, A5256801), 50 µg/mL gentamicin (Gibco, Thermo Fisher Scientific, 15710064), 25 µg/mL bovine fetuin (Sigma-Aldrich, F3004), 5 ng/mL recombinant human EGF (Thermo Fisher Scientific, PHG0311), 0.5 ng/mL recombinant human bFGF (Thermo Fisher Scientific, PHG0026), 2.5 ng/mL recombinant human HGF (Sino Biological, 10463-HNAS), 0.2 µg/mL dexamethasone (Sigma-Aldrich, D4902), and 5 µg/mL recombinant human insulin (Sigma-Aldrich, 91077C). Cells were maintained at 37°C in a humidified atmosphere of 5% CO₂. Upon reaching approximately 80% confluence, myoblasts were seeded at 3 × 10⁵ cells per well into six-well plates pre-coated with 0.5% gelatine. After 24 hours, growth medium was replaced with serum-free co-culture differentiation medium consisting of DMEM supplemented with 10 µg/mL insulin and 50 µg/mL gentamicin. GFP-labelled NPCs were added at a final concentration of 2.5 × 10⁴ cells/mL, and co-cultures were maintained for up to 12 days under humidified 5% CO₂ conditions with medium replaced every 48 hours. Functional NMJ formation was assessed daily by phase-contrast microscopy. Successful co-cultures exhibited progressive motor neuron neurite extension, direct contact between GFP-positive axons and differentiated myotubes, and spontaneous myotube contractions typically emerging by day 7. Cultures were used for downstream analyses on day 12, when NMJ formation was established structurally and functionally.

**Supplementary Methods S2. Model Validation**

The following experiments were performed to confirm motor neuron identity, structural NMJ formation, and functional synaptic connectivity in the co-culture system prior to AGE treatment. Motor neuron identity within the co-culture was confirmed by immunostaining for βIII-tubulin. Cells were fixed, permeabilised, and blocked as described in the main Materials and Methods. Primary antibody: (anti-βIII-tubulin; 1:400, eBioscience, AB_2574421). Structural NMJ formation was confirmed by co-staining with α-bungarotoxin conjugated to Alexa Fluor™ 647 (1 µg/mL; Invitrogen, B35450) applied for 30 min at room temperature prior to fixation, labelling acetylcholine receptor clusters at sites of motor neuron myotube contact.

To pharmacologically validate functional synaptic connectivity, spontaneous contraction recordings were performed immediately before and after application of α-bungarotoxin (1 µg/mL; Invitrogen, B35450) to established day 12 co-cultures. α-Bungarotoxin is a neurotoxic peptide derived from the venom of the snake Bungarus multicinctus that irreversibly binds to postsynaptic nicotinic acetylcholine receptors at the NMJ, preventing acetylcholine binding and abolishing neuromuscular transmission. Contraction intensity was monitored at multiple time points over a 30-minute recording window and further assessed at 1.5 and 24 hours following medium replacement. The complete and sustained abolition of spontaneous contractions confirmed that contractile activity in the co-culture model depends on AChR signalling at the NMJ. In separate experiments, L-glutamic acid (400 µM; Sigma-Aldrich, G1251) was applied to day 12 co-cultures. L-glutamic acid acts on motor neurons rather than directly on myotubes, stimulating depolarisation and acetylcholine release into the synaptic cleft, which subsequently activates postsynaptic AChRs to enhance myotube contractile activity. Contraction intensity was recorded at multiple time points over 30 minutes following application and at 1.5 and 24 hours after medium replacement, confirming functional motor neuron excitation and downstream myotube activation. Contraction analysis was performed as described in Supplementary Methods S3.

**Supplementary Methods S3. Quantitative Analysis of Myotube Contraction**

To assess functional neuromuscular connectivity in the co-culture system, myotube contraction intensity was quantified using video microscopy. On day 12 of co-culture, when NMJ formation and contractile activity had been established, live phase-contrast videos were captured from ten randomly selected fields of view using a Carl Zeiss Axio Observer 7 phase-contrast microscope at 20× magnification. Co-culture plates were maintained on an inverted microscope stage within an incubation chamber at 37°C and 5% CO₂ throughout the recording process. Following a few minutes of live observation to confirm consistent contractile activity, real-time video recordings lasting 35 seconds (940 frames) were captured at 27 frames per second. For data analysis, MATLAB (MathWorks R2022a, version 9.12.0) was used with an in-house motion analysis algorithm. Contraction intensity was quantified through frame-by-frame analysis of pixel intensity changes; for each consecutive frame pair, the absolute difference in pixel intensity values was calculated, summed across all frames, and normalised to the total number of frames, yielding a mean contraction intensity expressed in arbitrary units (a.u.). Spontaneous contraction frequency was calculated as the total number of contractile events divided by the total recording duration, expressed as contractions per second (s⁻¹). An illustrative video of the contraction analysis method is available at:

[Myotube Contraction Analysis](https://mmutube.mmu.ac.uk/media/Myotube+Contraction+Analysis/1_fee4fyqs)
